# 3D Bioprinting of Kidney Tissue Using a Photocrosslinkable Hydrogel Derived from Decellularized Extracellular Matrix

**DOI:** 10.1101/2025.02.21.639595

**Authors:** Jaemyung Shin, Nima Tabatabaei Rezaei, Subin Choi, Keekyoung Kim

## Abstract

Three-dimensional bioprinting has emerged as a promising strategy in tissue engineering, aiming to fabricate functional tissue constructs for organ regeneration. A critical challenge in this field is the development of organ-specific bioinks that can provide a microenvironment conducive to cellular growth and differentiation. In this study, we successfully developed a photocrosslinkable bioink by methacrylating decellularized porcine kidney extracellular matrix. The decellularization process effectively removed all cellular components while preserving the native kidney extracellular matrix composition. The resulting methacrylated decellularized extracellular matrix bioink exhibited optimal rheological properties, making it well-suited for digital light processing based stereolithography and piston-driven extrusion bioprinting. Human embryonic kidney cells encapsulated in the bioink showed high viability and a strong proliferative capacity, indicating potential for tissue-specific maturation over time. This work demonstrates the feasibility of utilizing kidney-specific decellularized extracellular matrix-based bioinks, providing a platform for engineering renal tissue constructs for therapeutic applications.

## 1. Introduction

Chronic kidney disease (CKD), which can progress to end-stage renal disease, is a significant health concern in North America, affecting approximately 37 million people in the United States, accounting for about 15% of the adult population [1],[2]. This condition is a leading cause of morbidity and mortality, contributing to over 100,000 new cases of end-stage renal disease annually [3]. In Canada, CKD affects over 3 million individuals, with a prevalence rate of around 10% among adults [4]. Given the severity and prevalence of kidney disease, bioengineering approaches, such as tissue bioprinting, offer promising solutions to address these challenges.

Three-dimensional (3D) bioprinting technology provides significant advantages for biomedical engineering applications, particularly in the development of personalized tissue models, organ prototypes, and patient-specific implants [5]. Its ability to fabricate complex tissue structures with high precision allows for better replication of native tissues, which is crucial for applications such as tissue engineering and precision medicine [6]. One of the most critical components of bioprinting is bioink, which must meet essential criteria, including biocompatibility, printability, and biodegradability to ensure successful bioprinting and functionality of the bioprinted tissues or organs [7], [8].

Bioinks are commonly derived from both synthetic and naturally occurring hydrogels. Synthetic biomaterials offer advantages such as customizability, consistency, and scalability, allowing for precise property control and large-scale production [9]. However, naturally derived biomaterials, particularly those based on extracellular matrix (ECM), are valued for their biocompatibility and ability to closely replicate the native tissue microenvironment, which is essential for promoting cellular growth and tissue regeneration [10]. Biological scaffold materials composed of ECM have been shown to promote structural remodeling of various tissues in both preclinical animal studies and human clinical applications [11].

In this study, we developed a kidney ECM-based hydrogel precursor that retains its natural biochemical and structural properties. However, decellularized ECM (dECM), in particular, has poor mechanical properties, making it challenging to use for printing applications [12]. ECM-based hydrogels are often limited by their slow and uncontrollable gelation processes and mechanical properties that fail to replicate physiological conditions. To address these challenges, natural polymers were functionalized with photoreactive components to enable rapid formation of hydrogels with tunable gelation properties. Synthetic modifications, including photocrosslinkability, were also incorporated to enhance functionality and mechanical properties. Tabatabaei Rezaei *et al.* developed a methacrylated dECM (dMA)-based bioink from porcine liver and formulated a hybrid bioink by incorporating it with gelatin methacrylate (GelMA). The photocrosslinkable dECM-based bioink preserved the beneficial properties of dECM, as demonstrated by a significant increase in HepG2 cell proliferation and cluster formation, compared to dECM- free bioinks. However, using highly substituted dMA as a standalone bioink offers significant advantages over hybrid formulations. With a high degree of substitution (DS) of methacrylate group, dMA undergoes solely photocrosslinking, eliminating the need for additional polymers like GelMA while maintaining structural integrity. Moreover, the absence of non-native components reduces the risk of immune responses, making pure dMA an ideal bioink for fabricating biomimetic hepatic tissues. In this study, we developed a kidney dECM methacrylate (KdMA) bioink for digital light processing-based stereolithography (DLP-SLA) and piston-driven extrusion bioprinting without the addition of supplementary materials. Additionally, we aimed to evaluate whether the KdMA hydrogel could promote tissue formation of human embryonic kidney cells at a level comparable to the commonly used, well-established GelMA bioink [13].

Previous studies, such as the work by Visscher *et al.* have demonstrated the viability of kidney ECM-derived bioinks for extrusion-based bioprinting, showing successful cell proliferation within printed constructs [14]. However, our approach introduces several significant advancements and unique characteristics that differentiate it from existing methods. First, we developed a cost-effective and faster immersion-based decellularization process compared to the perfusion method. Then, this dMA bioink was used to fabricate a 3D kidney scaffold via the DLP-SLA bioprinting method. This printing method enabled higher-resolution printing through layer-by-layer crosslinking of the bioink using 405 nm light, achieving faster printing speeds. Also, we printed the KdMA bioink using the custom-modified extrusion bioprinter to verify its printability and cellular biocompatibility. Using KdMA, we successfully promoted cellular regeneration in renal tissue bioprinting [15].

As the kidney is considered a soft organ with a shear modulus of approximately 4.5 kPa, kidney organoids naturally prefer to grow in soft microenvironments that better mimic the native ECM [16]–[18]. Nerger e*t al.* encapsulated stem cell-derived kidney organoids on day 7 of differentiation in Matrigel, Collagen Type I, and alginate hydrogel, and conducted a comparative analysis. The results demonstrated that organoids encapsulated in alginate hydrogel exhibited stable differentiation without significant cell migration, with all major nephron segments remaining present even after 21 days of differentiation [19]. They also demonstrated that kidney organoids exhibited improved growth and development in a soft and viscoelastic hydrogel, highlighting the importance of ECM mechanics in organoid maturation. Therefore, our next study aims to encapsulate induced pluripotent stem cell (iPSC)-derived kidney organoids within a KdMA bioink to further enhance maturation and angiogenesis, creating a more physiologically relevant microenvironment. The bioink developed in this study not only addresses the poor printability of dECM, improving its printability but also offers a versatile microenvironment by allowing precise control of its mechanical properties through simple adjustment of light exposure and material concentration. This adaptability will be highly advantageous for the printing of kidney organoids. Furthermore, by encapsulating renal cells in the developed bioink and culturing them for extended periods, a more realistic drug screening platform can be created compared to traditional 2D culture methods. This highlights the bioink’s broad versatility and significant potential for various applications.

## 2. Materials and methods

### 2.1. Decellularization of porcine kidney

The kidney dECM was developed through modifications and further optimization of previously reported protocols [20]. Fresh porcine kidney tissue was purchased on the day of slaughter from a local butcher, transported on ice to the laboratory, and processed by removing the renal capsule and perirenal fat before freezing. The kidney tissue was then stored at -20°C to facilitate consistent slicing. The following day, the tissue was sliced into 0.1–0.3 cm thick sections using a meat slicer. To remove blood, the sliced kidney tissue was washed three times for 30 minutes each with distilled water at a volume ten times greater than the tissue volume in a 3500 mL beaker. The tissue was then treated with 0.5% Triton X-100 in 1M NaCl for 16 hours. The next day, the tissue was rinsed three times for 1 hour each with distilled water.

Following this, the tissue was stirred in a DNase solution at 37°C for 6–7 hours, and then washed with phosphate buffered saline (PBS) for an additional 12 hours. Dnase is an important enzymatic agents that cleave nucleic acid sequences, facilitating the removal of nucleotides after cell lysis in tissues. The following day, the tissue was treated with a 0.1% peracetic acid solution for 1 hour under agitation, followed by three washes with distilled water, each lasting 30 minutes. At this stage, the kidney tissue, initially pink, had turned transparent. The decellularized kidney tissue was then frozen at -20°C for one day and freeze-dried for three days. The kidney dECM, finely ground to achieve an ultra-small particle size, was dissolved at a concentration of 1 mg/mL in 0.5 M acetic acid containing pepsin, and the mixture was incubated at room temperature for 48 hours.

To induce precipitation, sodium chloride was added to the solution to reach a final concentration of 5%. The solution was then centrifuged at 10,000 RPM for 15 minutes, and the precipitated tissue was collected. The recovered kidney dECM was subjected to dialysis using 3.5 kDa tubing at 4°C, with two solution changes per day, followed by freeze-drying to complete the solubilization process. The detailed description of the methacrylation process is provided in section 2.3.

### 2.2. Histological characterization and staining analysis of decellularized extracellular matrix

To visualize and analyze residual porcine cells and microarchitecture in tissues following the decellularization process, both untreated native kidney tissue and kidney dECM were fixed in 4% (wt/vol) paraformaldehyde solution at room temperature for 24 hours. The samples were then dehydrated by sequential immersion in ethanol solutions of 100%, 95%, 90%, 75%, 70%, and 50% (vol/vol) overnight. Subsequently, the tissues were embedded in paraffin. For further staining procedures, the paraffin-embedded tissue samples were sectioned into thin slices with a thickness of 5–7 μm. These sections were stained with hematoxylin and eosin (H&E) (VWR, Mississauga, ON, Canada) and Masson’s trichrome stain (Polysciences Inc., Warrington, PA, USA). Staining enabled the detection of any residual cellular components and provided insight into the preservation of structural integrity in the decellularized tissue. This approach ensured a comprehensive evaluation of the effectiveness of the decellularization process and the structural characteristics of the tissue. The samples of the H&E and Masson’s trichrome were further subjected to DAPI staining and imaged using an inverted microscope (ECHO Revolve, San Diego, CA, USA). This process enabled the visualization of nuclei within the kidney tissue and was used to evaluate the presence or absence of nuclei in the dECM samples.

The quantification of DNA and total protein concentration (TPC) was performed using the PicoGreen dsDNA Assay Kit (ThermoFisher, Waltham, Massachusetts, USA) and the TPC kit (ThermoFisher, Waltham, Massachusetts, USA), respectively, to evaluate the residual cellular and protein content in dECM (*n*=3). For this analysis, 50 mg of native kidney tissue and lyophilized kidney dECM were added to 1 mL of a pepsin digest solution (0.1 mg/mL, Ward’s Science, ON, Canada). The mixture was homogenized using an FSH- 2A High-Speed Homogenizer at 20,000 RPM. Subsequently, the samples were vortexed and incubated in a 65°C water bath for 6 hours. After incubation, the samples were centrifuged at 4,000 RPM for 15 minutes, and the supernatant was collected. Measurements were performed using a microplate reader (SpectraMax® M3, Molecular Devices, San Jose, CA, USA) according to the manufacturer’s protocol.

### 2.3. Methacrylation of decellularization of extracellular matrix

The methacrylation process for the solubilized dECM was conducted as follows: 1 g of kidney dECM was dissolved in 100 mL of 0.5 M acetic acid at room temperature. To optimize methacrylation activation, 5 M NaOH was added to adjust the pH of the solution to 8–9. Methacrylic anhydride (Sigma-Aldrich, St. Louis, MO, USA) was then introduced at a concentration of 2.5 mL per gram of dECM, added dropwise to ensure controlled reaction conditions. The solution was stirred continuously at room temperature for 2 days to facilitate synthesis. Following this, the reaction product was subjected to dialysis using a 3.5 kDa dialysis tube (Thermo Fisher Scientific, Waltham, MA, USA) at 4°C for three days, with water replaced twice daily. The final sample was freeze-dried for three days to obtain the final KdMA.

The DS of the synthesized KdMA was quantified using proton nuclear magnetic resonance (^1^H NMR) spectroscopy, ensuring the efficiency and consistency of the methacrylation process. The KdMA solution was prepared at a concentration of 0.5% (w/v) 1 mL of deuterium oxide (ThermoFisher, Waltham, Massachusetts, USA). Next, ^1^H NMR spectra for the synthesized KdMA samples were recorded using a 600 MHz NMR spectrometer. The internal reference was set to hydroxyl signals (0.5-1 ppm). The peaks corresponding to primary amine groups (2.8-2.95 ppm) were integrated to calcualte the DS.

### 2.4. Preparation of precursor hydrogel and bioprinting system

Three different concentrations of KdMA were dissolved in sterilized PBS. To prepare these solutions, a 0.5% (w/v) stock solution of lithium phenyl-2,4,6- trimethylbenzoylphosphinate (LAP) (Sigma-Aldrich, St. Louis, MO, USA), was first prepared. Varying amounts of KdMA, corresponding to different weights of the compound, were then added to the stock solution, and the mixture was stirred at room temperature overnight. To ensure uniformity, stirring was continued for up to three days. Mixing for less than three days resulted in the presence of fine fibers that hindered crosslinking, while stirring for more than a week was found to decrease the crosslinking ability. The LAP photoinitiator consists of lithium ions and an acyl phosphinate group, which combine to be activated by ultraviolet (UV) or visible light (particularly at the wavelength of 405 nm) to initiate polymerization. Due to its high photoinitiation efficiency and low toxicity, LAP is frequently used in biomaterials research and applications. The exposure duration was varied for each bioink combination based on its photocrosslinking characteristics.

### 2.5. Characterization of mechanical, swelling, and degradation properties of the developed precursor hydrogel

The mechanical properties of the hydrogel were evaluated through a comparative analysis of the compressive modulus of crosslinked hydrogels. Compression testing was conducted using a flat cylinder probe with a diameter of 12.5 mm. To prepare the samples, 2 mL of hydrogel precursor solution with varying KdMA concentrations was transferred into cylindrical molds with a diameter of 8 mm and a height of 4 mm (**Figure S1**). Each well received 400 µL of prepolymer solution, which was crosslinked for 30 seconds in the DLP-SLA bioprinting system. The shape of the samples used for mechanical testing was different from those used in rheological measurements. In rheological measurements, the samples were thin, exposing a larger surface area to light, whereas the mechanical samples were cylindrical, with a smaller surface area exposed to light, requiring a longer crosslinking time. Compression was applied to the hydrogel surface up to 80% strain, and force-displacement data were recorded. The compressive modulus was determined using the initial dimensions of the hydrogel samples and a custom MATLAB script, with the slope of the linear region corresponding to an initial strain of 10%.

The swelling ratio of the crosslinked hydrogel samples was measured to evaluate their water uptake capability. A total of 400 µL of hydrogel solutions with different concentrations were cast into molds with a diameter of 8 mm and a height of 4 mm and crosslinked using a DLP-SLA bioprinter. The crosslinked hydrogels were then immersed in PBS and stored at 37°C with 5% CO_2_. At predetermined time points, the samples were removed and the hydrated weight (W_w_) of each sample was recorded. Subsequently, the samples were frozen at -20°C and lyophilized for three days to obtain the dry weight (W_d_). The swelling ratio was then calculated using the following equation:

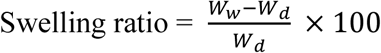

To investigate the degradation behavior of the samples, hydrogels were prepared using the previously described method. All samples were freeze-dried for three days, and their initial dry weight (W_o_) was recorded. The samples were then immersed in PBS at 37°C. At predetermined time points, samples were retrieved for degradation analysis. The retrieved samples were frozen at -20°C and subsequently freeze-dried for three days. The weight of the freeze-dried samples was then measured to determine the remaining polymer matrix weight after degradation (W_r_). The degradation ratio was expressed as the percentage of the remaining weight of the sample after degradation. The equation used for the calculation is as follows:

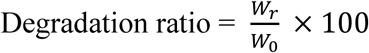

### 2.6. Scanning electron microscopy for porosity quantification

The microstructure of the hydrogels was meticulously examined using scanning electron microscopy (SEM). Hydrogel samples were prepared according to previously established protocols, followed by freezing at -20°C and lyophilization for three days. To capture naturally fractured cross-sections, the freeze-dried samples were partially sectioned with a sharp blade, allowing the remaining portions to fracture spontaneously. The fractured surfaces were positioned with the exposed surfaces facing upwards and carefully mounted onto SEM stubs to ensure proper alignment for imaging. Before imaging, the samples were sputter-coated with a 15 nm thin layer of gold to enhance conductivity and minimize charging effects. High-resolution imaging was then performed using a Phenom Pro X SEM, allowing for detailed visualization and analysis of the microstructural characteristics of the hydrogels.

### 2.7. Characterization of crosslinking kinetics and rheological properties

The rheological properties of KdMA at varying concentrations were meticulously characterized using a rheometer (MCR 302, Anton Paar, Graz, Austria). All measurements were performed at ambient temperature employing a parallel plate geometry with a 25 mm diameter and a 0.5 mm gap maintained on the rheometer platform. To evaluate photocrosslinking kinetics, a 405 nm light source was applied from beneath to irradiate a thin prepolymer layer, initiating the crosslinking process. The light exposure began 60 seconds after the experiment commenced and was maintained for a duration of five minutes. Oscillatory measurements were conducted at a constant 0.5% shear strain and a frequency of 1 Hz, allowing for the simultaneous recording of both storage and loss moduli. After the light exposure phase, frequency sweeps were performed within the range of 0.1–100 rad/s while maintaining a constant shear strain of 0.5%. This was done to observe the dynamic rheological properties of the bioink as it underwent crosslinking.

To validate the shear-thinning behavior of the bioink, viscosity measurements were performed by varying the shear rate from 0.01 to 1000 s⁻¹ in the absence of light exposure. Additionally, the three-interval thixotropy test was employed to assess the thixotropic behavior and phase transitions following exposure to excessive stress. During the first (0 to 60 seconds) and third (91 to 270 seconds) intervals, low shear stress (0.1 Pa) was applied to stabilize the sample and allow for structural recovery. Conversely, high shear stress was applied during the second interval (61 to 90 seconds) to simulate the flow through the nozzle and the structural deformations that occur during extrusion bioprinting.

### 2.8. Cell culture

Human embryonic kidney (HEK) cells were maintained in high-glucose Dulbecco’s Modified Eagle Medium (Corning, Arizona, USA) supplemented with 10% fetal bovine serum (Corning, Arizona, USA) and 1% penicillin/streptomycin (Cytiva, Marlborough, MA, USA). The cells were cultured in tissue culture flasks and passaged as necessary, with growth conditions sustained in a 37°C incubator with 5% CO_2_. The culture medium was refreshed every two to three days to support optimal cell viability and proliferation.

### 2.9. Characterization of cell viability in 3D-printed hydrogel constructs

To fabricate cell-encapsulated scaffolds, cells were seeded at a density of 1×10^6^ cells/mL within various bioinks. Following crosslinking, the scaffolds were cultured in well plates, with medium changes performed every three days. Cell viability was assessed at the desired time point post-bioprinting using the LIVE/DEAD Viability/Cytotoxicity Kit (Biotium, Fremont, CA, USA). To ensure thorough removal of residual culture medium, the 3D scaffolds were washed three times with PBS, followed by a 15-minute incubation in PBS at room temperature. This additional incubation step was essential as residual medium often remains trapped within the porous structure of 3D scaffolds despite washing. Extended exposure to PBS facilitates the diffusion and removal of any remaining medium, ensuring more consistent and uniform staining.

For viability assessment, the cell-laden scaffolds were stained using a solution of 0.5 μL/mL calcein AM and 2 μL/mL ethidium homodimer (EthD) dissolved in PBS. The staining solution was incubated in the dark at room temperature for 30 minutes. After staining, scaffolds were washed three times with PBS to eliminate any unbound reagents. Fluorescence imaging was then performed using an inverted fluorescence microscope to visualize live (calcein AM) and dead (EthD) cells, as well as the overall distribution of cells within the scaffolds. Calcein AM fluorescence was detected using the FITC channel, while EthD fluorescence was visualized using the TxRed channel. High-resolution 3D imaging and z-stack reconstruction of the cell-encapsulated hydrogels were carried out using a Nikon Eclipse Ti confocal microscope (Nikon, Tokyo, Japan).

### 2.10. Quantitative analysis of cell proliferation via XTT assay

To evaluate cellular proliferation and assess mitochondrial metabolic activity, the tetrazolium hydroxide salt assay (XTT kit, Biotium, Fremont, CA, USA) was employed. The XTT reagent was added to the culture medium at a 1:10 ratio to enable the measurement of cellular viability and metabolic activity. The samples were incubated with the XTT solution for 24 hours at 37°C in a 5% CO_2_ atmosphere to allow for sufficient reduction of the tetrazolium salt by metabolically active cells. Following incubation, 100 μL of the supernatant from each well was carefully transferred to a 96-well plate to minimize potential interference from cellular debris. The metabolic activity of the cells was then quantified by measuring the absorbance of the reduced formazan product, which forms as a result of the mitochondrial enzymatic reduction of the XTT reagent. Absorbance readings were taken at a wavelength of 450 nm using a spectral scanning plate reader, with the absorbance intensity correlating with the number of viable cells and their metabolic activity. This method provides a reliable and quantitative assessment of cellular proliferation and mitochondrial function in response to biomaterial or experimental conditions.

### 2.11. Assessment of cell proliferation and morphology in bioprinted scaffold

To evaluate cell proliferation within scaffolds fabricated from the KdMA bioink, HEK cells were encapsulated and cultured for one month. The bioink, seeded at a density of 1×10^6^ cells/mL, was loaded into a syringe mounted on an extrusion-based bioprinter, and an appropriate volume of bioink was dispensed. Multiple samples with a diameter of 2 cm and a thickness of 200 μm were printed to ensure that at least one dimension fell within a range comparable to the critical limit for medium diffusion. Immediately after printing, crosslinking was performed using 405 nm light. The crosslinked discs were transferred to a petri dish and thoroughly washed multiple times with PBS to remove any uncrosslinked bioink. Finally, the samples were transferred to an incubator and cultured for one month, with the culture medium replaced with fresh medium every two to three days.

To evaluate cell morphology and proliferation, at least three smaller discs, approximately 4 mm in diameter, were created from each sample using a disc punch. The samples were stained for cytoskeleton and nuclear visualization at 5, 10, and 30 days of culture using Phalloidin (Cytoskeleton Inc., Denver, CO, USA) and DAPI (MilliporeSigma, Oakville, ON, Canada), respectively. Before staining, the samples were fixed in 4% v/v paraformaldehyde for 90 minutes. Permeabilization was achieved using 0.5% v/v Triton X-100 for 20 minutes to enhance membrane permeability, facilitating the penetration of staining reagents into the cells. The samples were then stained in the dark at room temperature for 90 minutes with Phalloidin 488. Following staining, the samples were mounted using a mounting medium and imaged using a fluorescence microscope equipped with DAPI and FITC channels.

### 2.12. Statistical analysis

Quantitative data are reported as mean ± standard deviation and inferential statistics (p-values) were used for further analysis. Statistical significance was determined using one-way ANOVA, with significance levels set at p < 0.05, p < 0.01, or p < 0.001. To identify specific differences between group means, Tukey’s HSD test was used for post hoc analysis. The analysis was performed using GraphPad Prism version 12.2.2 software.

## 3. Results and discussion

### 3.1. Assessment of decellularization efficiency of kidney tissues

To develop a kidney dECM bioink suitable for DLP-SLA bioprinting, a decellularization process was performed using a fresh porcine kidney (**Figure 1A**). The process was optimized to ensure the complete removal of porcine cells while preserving essential kidney ECM components, such as collagen fibers. As shown in **Figure 1B**, the kidney dECM exhibited thermocrosslinking behavior at 37°C, attributed to its collagen content. To further investigate its composition, histochemical staining was performed to visualize cellular nuclear material and collagen content. **Figure 1C** displays representative cross-sectional images of native kidney tissue, stained with H&E, highlighting well-defined nephron structures, including glomeruli, and organized cellular arrangements within the kidney cortex, thereby reflecting the intact architecture of the kidney. In contrast, H&E staining of the kidney dECM sample shows a structure lacking the dark purple-stained nuclei observed in the native kidney tissue, confirming successful cell removal. Additionally, the presence of collagen in the kidney dECM was verified through Masson’s trichrome staining, which highlighted collagen fibers in blue. This result confirms that collagen fibers remain the primary structural component following decellularization.

**Figure 1.**
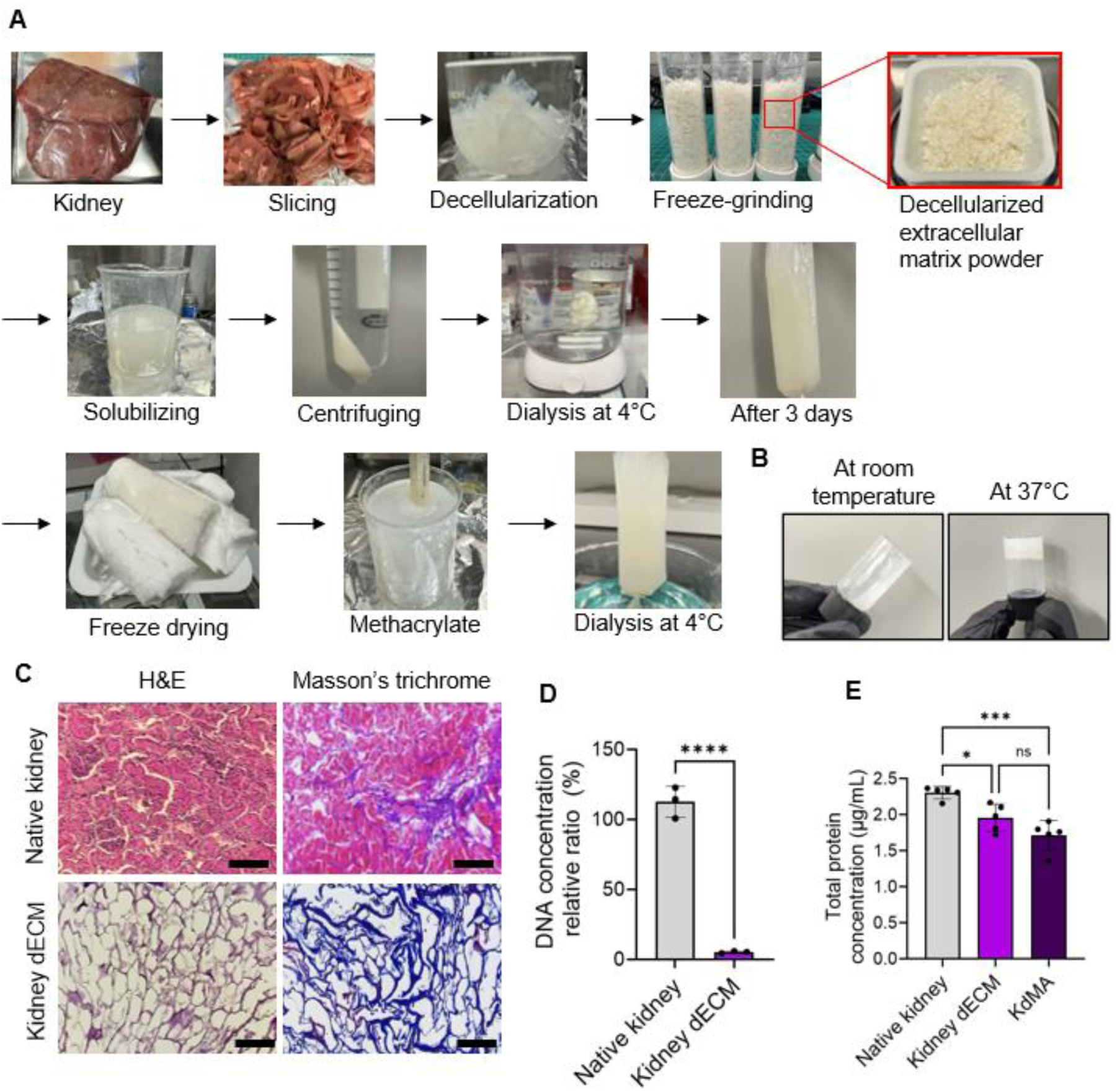
(A) A comprehensive overview of the process for synthesizing KdMA from porcine kidney tissue. (B) Thermal gelation behavior of kidney dECM hydrogel demonstrating the transition from sol to gel at physiological temperature. (C) Hematoxylin and eosin staining and Masson’s trichrome staining of the native kidney and KdMA. Scale bars, 100 μm. (D) Comparison of total DNA content between the native kidney and kidney dECM (*n*=5; ** p < 0.01). (E) Comparative analysis of total protein concentration among the native kidney, kidney dECM, and KdMA. (*n*=5; * p < 0.05, *** p < 0.001)

To quantitatively assess the efficiency of decellularization, DNA and protein content were analyzed. The results showed that the DNA content of the kidney dECM samples decreased to less than 5.21% of the native kidney DNA content (**Figure 1D**). Although ionic treatments such as sodium dodecyl sulfate (SDS) are highly effective in cell removal, they are more aggressive than non-ionic treatments and tend to wash away a greater proportion of ECM proteins. To evaluate the impact of the washing and functionalization step on TPC, measurements revealed

### 3.2. Functionalization and characterization of KdMA bioink

The abundant collagen protein within the synthesized kidney dECM was utilized to modify the amine (-NH_2_) functional groups in the primary collagen framework, allowing for the substitution with methacrylate groups (**Figure 2A**). Although kidney dECM was methacrylated using methacrylic anhydride to synthesize KdMA, **Figure 2A** depicts the GelMA synthesis process, where gelatin is methacrylated with methacrylic anhydride. This illustration is included to help understanding, as the methacrylation process for both GelMA and KdMA follows a similar chemical reaction mechanism. The methacrylation process of collagen requires dissolution; thus, pepsin was used to enzymatically digest the matrix, partially hydrolyzing non-fibrillar collagen regions to facilitate solubilization while preserving its triple-helical structure, which is essential for maintaining bioactivity. This enzymatic digestion also ensured a homogeneous solution, enhancing reactivity with methacrylic anhydride for efficient methacrylation.

**Figure 2.**
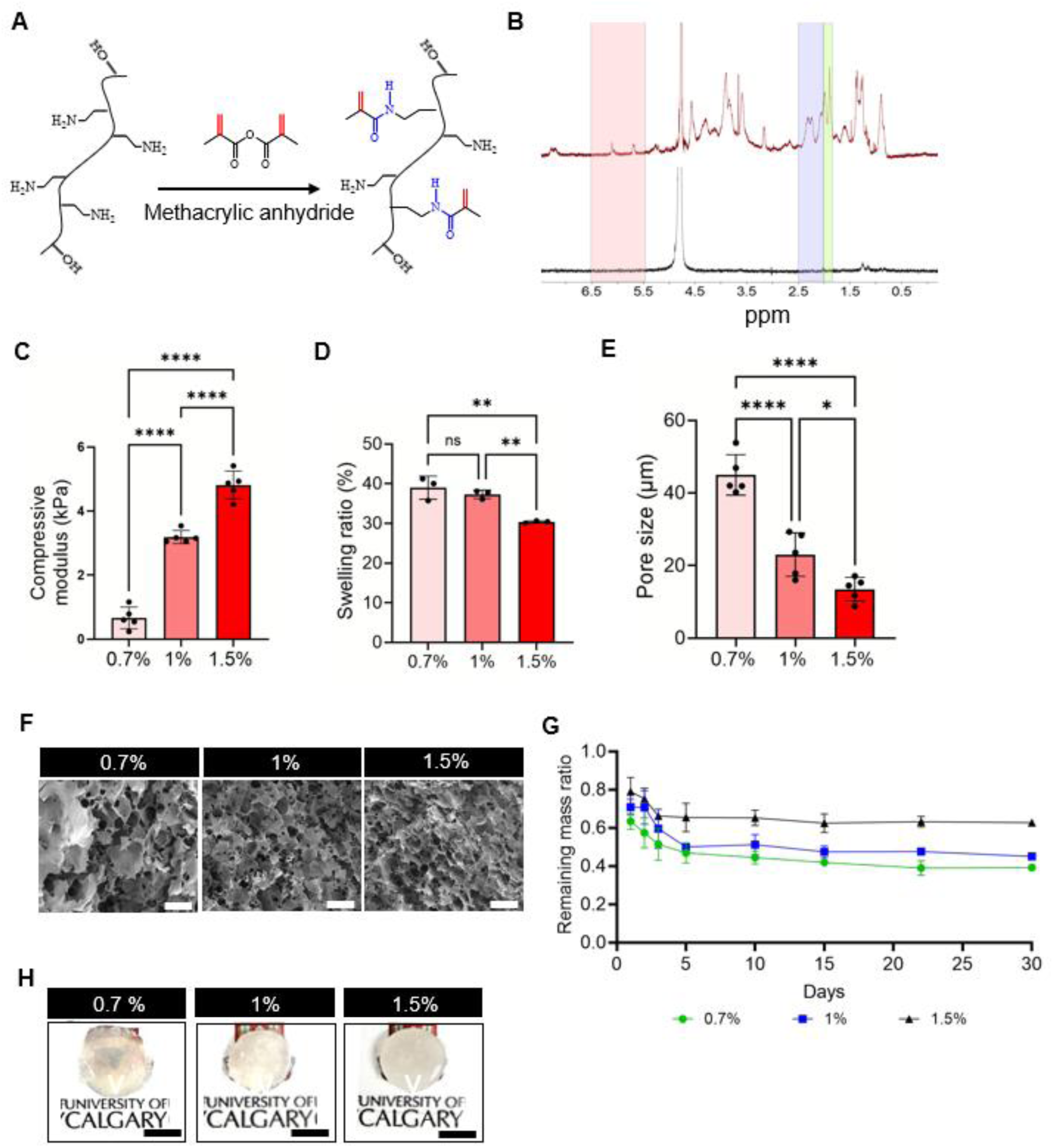
(A) Mechanism of methacrylate group substitution on KdMA backbone molecules. (B) Characterization of the DS of methacrylate group in KdMA, as demonstrated by ^1^H NMR spectra of KdMA and kidney dECM samples. (C) Compressive modulus of hydrogels with different compositions (*n*=5). (D) Mass swelling ratio of hydrogels with different compositions (*n*=5). (E) Pore size analysis of the 3D microstructure of the hydrogels (*n*=5). (F) Scanning electron microscopy images showing the porous structure. Scale bars, 200 μm. (G) Degradation analysis of the hydrogels immersed in sterilized PBS (*n*=3). (H) Transparency of disk-shaped hydrogels with different compositions of KdMA. Scale bars, 5 mm.

To assess the DS of methacrylate groups, ^1^H NMR analysis was performed (**Figure 2B**). The KdMA exhibited new peaks at 5.29 and 5.72 ppm, corresponding to the methacrylate end groups, and at 1.97 ppm, corresponding to the methacrylamide end groups of the macromolecules. Peak integration relative to the internal reference revealed a DS of approximately 61.5% [11],[12]. To further enhance the DS, an additional grinding step using a cryomilling machine before the solubilization process mentioned in Section 2.1 is expected to improve methacrylation efficiency. Cryomilling reduces particle size and increases surface area, facilitating greater exposure of reactive sites and promoting more effective methacrylation. This process not only activates methacrylate groups but also improves material transparency by refining it into fine powders. As a result, it significantly decreases the stirring time required to achieve a homogeneous KdMA solution, which currently takes three days, ultimately enabling the production of a ready-to-use biomaterial with enhanced efficiency.

### 3.3. Measurements of mechanical, swelling, microstructure, and degradation properties

To investigate the relationship between KdMA concentration and compressive modulus, the compressive modulus of KdMA at different concentrations was measured. As the concentration increased from 0.7% to 1.5%, a significant increase in compressive modulus was observed (**Figure 2C**). The measured compressive moduli for 0.7%, 1%, and 1.5% KdMA were 0.67 kPa, 3.19 kPa, and 4.81 kPa, respectively. These results demonstrate that the compressive modulus can be increased by up to sevenfold compared to 0.7% KdMA. This tunability of mechanical properties makes the developed KdMA bioink a promising biomaterial for various kidney tissue modeling applications.

Material stiffness plays a critical role in mimicking the native kidney tissue environment [21]. Notably, nephron progenitor cells thrive in a soft environment with a stiffness range of 0.1–3 kPa, which supports kidney organoid formation [21]. Therefore, the development of a soft bioink was a key objective of this study. Various studies have explored the mechanical properties of healthy kidney tissue at different stages, as well as in diseased or cancerous kidneys [22]. Ruiter *et al.* investigated human iPSC-derived kidney organoids, designed to mimic human kidney organogenesis, under environments ranging from 0.1 to 20 kPa [21]. Organoids cultured in a 20 kPa hydrogel exhibited a significant reduction in interstitial and loop of Henle cells compared to those grown in 0.1 and 3 kPa environments. Additionally, organoids cultured in the 20 kPa hydrogel showed a marked decrease in lumen structures, along with the absence of key kidney cell types. These findings indicate that the stiffness and stress relaxation properties of the surrounding environment directly influence kidney organoid development, with soft hydrogels enhancing lumen structure maturation. The tunable flexibility of the KdMA hydrogel developed in this study, with adjustable stiffness ranging from 0.67 to 4.81 kPa, highlights its potential in promoting kidney organoid maturation. Furthermore, properly engineered extracellular matrices can be tailored to mimic pathological conditions, such as fibrosis, providing a platform for functional transplant development and advancing disease modeling applications.

The swelling ratio of hydrogels is a critical parameter in tissue engineering, affecting surface properties and solute diffusion [23]. Assessing the swelling ratio of crosslinked hydrogel samples provides insights into the degree of crosslinking and the material’s water absorption capacity [24]. As shown in **Figure 2D**, the swelling ratio decreases with increasing KdMA concentration. The 0.7% KdMA hydrogel exhibits the highest swelling ratio at 38.99%, while higher KdMA concentrations result in greater crosslinking density, reducing water retention capacity. Consequently, bioinks with higher KdMA concentrations demonstrate significantly lower swelling ratios; for instance, the swelling ratio of 1.5% KdMA is 30.35%, approximately 1.28 times lower than that of 0.7% KdMA. Interestingly, the difference in swelling ratio between 0.7% and 1% KdMA is not statistically significant, suggesting that the water retention capacity approaches a saturation point. Therefore, further reductions in KdMA concentration have a relatively minimal impact on the swelling ratio.

To further evaluate the morphology, porosity, and surface characteristics of the hydrogel, SEM was performed [25]. SEM analysis revealed that increasing KdMA concentration led to a decrease in pore size, which correlated with enhanced mechanical stiffness and a reduced swelling ratio (**Figure 2E**). As shown in **Figure 2F**, the 1.5% KdMA exhibited pores with an average size of 13.47 μm. This pore structure, as previously discussed, is associated with greater mechanical stiffness and lower swelling capacity. The 0.7% KdMA hydrogel exhibited larger pores measuring 44.98 μm, which correlated with the lowest compressive modulus and the highest swelling ratio.

Biodegradability is a highly desirable characteristic in tissue engineering applications, particularly for hydrogel materials, as it indicates how long the scaffold can maintain its structural integrity [26]. The developed bioink hydrogel exhibited a slow degradation rate, maintaining its structure for more than 30 days in PBS (**Figure 2G**). The 1.5% KdMA hydrogel exhibited the highest remaining mass ratio maintained over 30 days. This is likely attributed to the increased degree of crosslinking resulting from the higher concentration of methacrylate groups in KdMA. The elevated crosslink density necessitates the disruption of a greater number of covalent bonds during the degradation process, ultimately leading to structural collapse. Additionally, through expansion and microstructural evaluations, it was observed that as the KdMA concentration increased, pore size within the hydrogel decreased, resulting in a lower PBS absorption and a slower degradation rate. The ability to control the degradation rate of such hydrogels suggests their potential as versatile biomaterials for a wide range of applications, such as in vivo implantation and long-term in vitro studies [27]. **Figure 2H** illustrates the transparency of the crosslinked scaffold, revealing that its transparency is lower than expected. Moving forward, efforts should focus on incorporating cryomilling to further enhance the transparency of KdMA bioink.

### 3.4. Measurement of crosslinking time and rheological characterization

In DLP-SLA bioprinting, the successful application of kidney dECM-based bioinks depends on materials with precisely optimized rheological properties [28]. To ensure this, viscosity and crosslinking kinetics were assessed through rheological analysis. **Figure 3A** demonstrates the shear-thinning behavior of the developed bioink. Across all tested concentrations, the bioink exhibits a decreasing viscosity trend as the shear rate increases. This property is particularly critical for extrusion-based bioprinting processes. Under high shear conditions, such as during extrusion through a nozzle, the reduction in viscosity facilitates smooth deposition, while the subsequent recovery of viscosity ensures structural integrity post-printing. **Figure 3B** illustrates the relationship between shear stress and shear rate, which is essential for fluid model analysis. The slope of this graph corresponds to viscosity, and as the shear rate increases, the curve progressively flattens, further confirming the shear-thinning behavior of the bioink. Additionally, the graph does not start at zero shear stress; instead, flow begins only above a certain threshold (τ_y_), indicating that the bioink possesses a yield stress. This characteristic is crucial for maintaining structural stability after printing, as the presence of yield stress ensures that the bioink remains stationary below a specific stress threshold, thereby enhancing shape fidelity after bioprinting.

**Figure 3.**
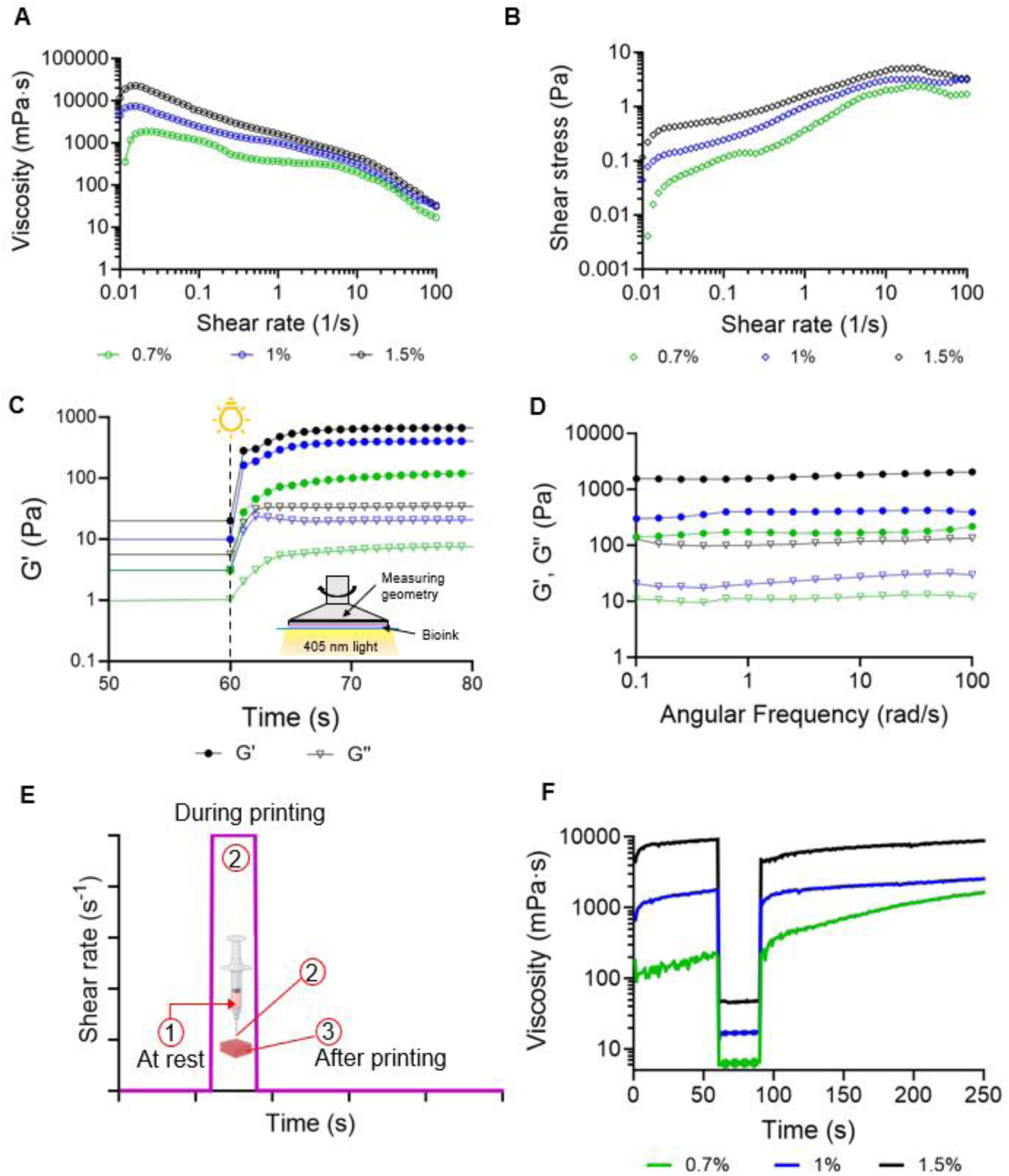
Dynamic rheological characterization of KdMA hydrogels. (A) Comparison of viscosity versus shear rate for different KdMA compositions. (B) Comparison of shear stress versus shear rate for different KdMA compositions. (C) Photocuring kinetics of KdMA hydrogels, represented by storage modulus (G’) and loss modulus (G’’) with 0.5% LAP. (D) Angular frequency response of bioink synthesized with 0.5% LAP. (E) Testing conditions of three-interval thixotropy test. (F) Three-internal thixotropy test results showing shear-thinning and structural behavior of the bioinks at different loading percentages.

As shown in **Figure S2**, bioinks with higher concentrations of KdMA exhibit higher viscosity, which is attributed to the greater content in KdMA, capable of undergoing methacrylate synthesis. According to the literature, the viscosity of resins optimal for SLA 3D printing should not exceed 5000 cP [29]. **Figure S2** shows that the viscosity of 0.7% KdMA was measured at 341.01 mPa.s, 1% KdMA at 1328.10 mPa.s, and 1.5% KdMA at 4494.27 mPa.s. Among these, the viscosity of 1.5% KdMA approaches the upper viscosity limit for SLA bioprinting, suggesting that the 1.5% KdMA bioink formulation may not be ideal for SLA bioprinting applications. However, the distinct shear thinning behavior exhibited by the 1.5% KdMA sample suggests its potential in various studies exploring extrusion-based 3D bioprinting using dECM-based bioink within the context of 3D bioprinting.

The photorheological properties of the bioink were investigated through photorheological testing, monitoring changes in the storage modulus (G’) and loss modulus (G’’) of various hybrid inks, as shown in **Figure 3C**. The changes in G’ and G’’ during the photopolymerization process provide valuable insights into several critical aspects of the material behavior. The point at which the G’ and G’’ curves intersect is typically considered the gel point of the material. Interestingly, for the KdMA samples, gelation does not occur within the light curing time. This observation suggests that the G’ exceeds G’’ prior to light exposure, likely due to the presence of collagen fibers, which form a partial network through entanglement before full light-induced gelation. This behavior appears to be more pronounced in higher-viscosity inks. Additionally, the gel time, defined as the period from the onset of light exposure to the observable increase in G’ and the initiation of gelation, varies among the samples. The 1.5%, 1%, and 0.7% KdMA bioinks all achieved full crosslinking within 10 seconds after the measurement began at 60 seconds of light exposure. However, the lowest concentration bioink exhibits a more gradual increase in storage modulus upon light exposure compared to the other two formulations. This indicates that the 0.7% KdMA bioink is the softest and undergoes a slower crosslinking process.

An important consideration in SLA printing is the ability to cleanly separate the printed structure from the uncured bioink, which is essential for achieving high printing resolution. Optimal printing requires high rigidity in the printed structure and low viscosity in the uncured bioink.ΔG’ is quantified as the logarithmic difference between the G’ values of the crosslinked and uncrosslinked states, serving to highlight the rheological disparity between the printed structure and the uncured hydrogel. Throughout the photocuring process, as the bioink undergoes increased crosslinking, the Δ G’ value rises until crosslinking is complete, at which point it stabilizes. However, variations in mechanical properties, as previously discussed, are also reflected in the different final G’ and G’’ values after crosslinking, which increase with rising KdMA concentration, thereby elevating Δ G’ as well. In conclusion, as the KdMA concentration increases, crosslinking time is shortened. This is due to the higher concentration of methacryloyl groups and the reduced free radial diffusion pathways, resulting in the formation of a stiffer structure.

In addition, the rheological stability of the photocrosslinked bioink was assessed using a frequency sweep test, where the storage modulus (G’) and loss modulus (G’’) were measured as a function of angular frequency (rad/s) (**Figure 3D**). As the values remained stable without significant fluctuations for up to five minutes, the graph is shown only up to 100 seconds for improved data visualization. All bioink formulations exhibited a consistent, linear trend without significant fluctuations, indicating that the bioink maintained a stable crosslinked network after photocuring. The absence of frequency dependence in G’ and G’’ suggests that the bioink achieved full crosslinking, forming a structurally stable hydrogel. This result confirms that the photocrosslinked bioink exhibits solid-like behavior with minimal viscoelastic variation across different angular frequencies, which is crucial for maintaining mechanical integrity after bioprinting.

To mimic bioprinting conditions, viscosity was measured as a function of time under three distinct phases: at rest, during printing, and after printing (**Figure 3E**). Before extrusion, the bioink remains in a static state within the syringe, maintaining high viscosity to prevent unwanted flow or leakage from the nozzle. Upon applied shear stress (e.g., nozzle extrusion), the bioink exhibits shear-thinning behavior, reducing viscosity to facilitate smooth deposition. This property is crucial for preventing nozzle clogging while ensuring uniform bioink flow. Finally, after deposition, the bioink undergoes viscosity recovery due to its thixotropic nature. Although the speed and extent of recovery determine the stability of the printed structure, rapid viscosity recovery ensures that the bioink does not spread or deform after printing, maintaining high resolution and structural fidelity. The relationship between shear rate and time was analyzed to further evaluate the thixotropic nature of the bioink (**Figure 3F**).

### 3.5. Assessment of the printing ability of the developed bioink

#### 3.5.1 Evaluation of the printability of the developed bioink in DLP-SLA bioprinting

The developed bioink was bioprinted using a customized DLP-SLA bioprinter, as shown in **Figure 4A**. To enable multilayer bioprinting, a z-stack movable plate was employed, with a Petri dish placed on top for layer-by-layer fabrication. Each bioprinted scaffold was fabricated with a single-layer volume of 100 µL, resulting in a total of 10 layers. The viability of encapsulated cells within the bioprinted scaffolds was assessed over 14 days. As shown in **Figure 4B**, cell viability remained above 90% from day 1 to day 14, indicating excellent cytocompatibility of both the bioink and the DLP-SLA bioprinting process. These results suggest that the photopolymerization process did not exert significant cytotoxic effects and that the bioprinted microenvironment successfully supported cell survival and proliferation.

**Figure 4.**
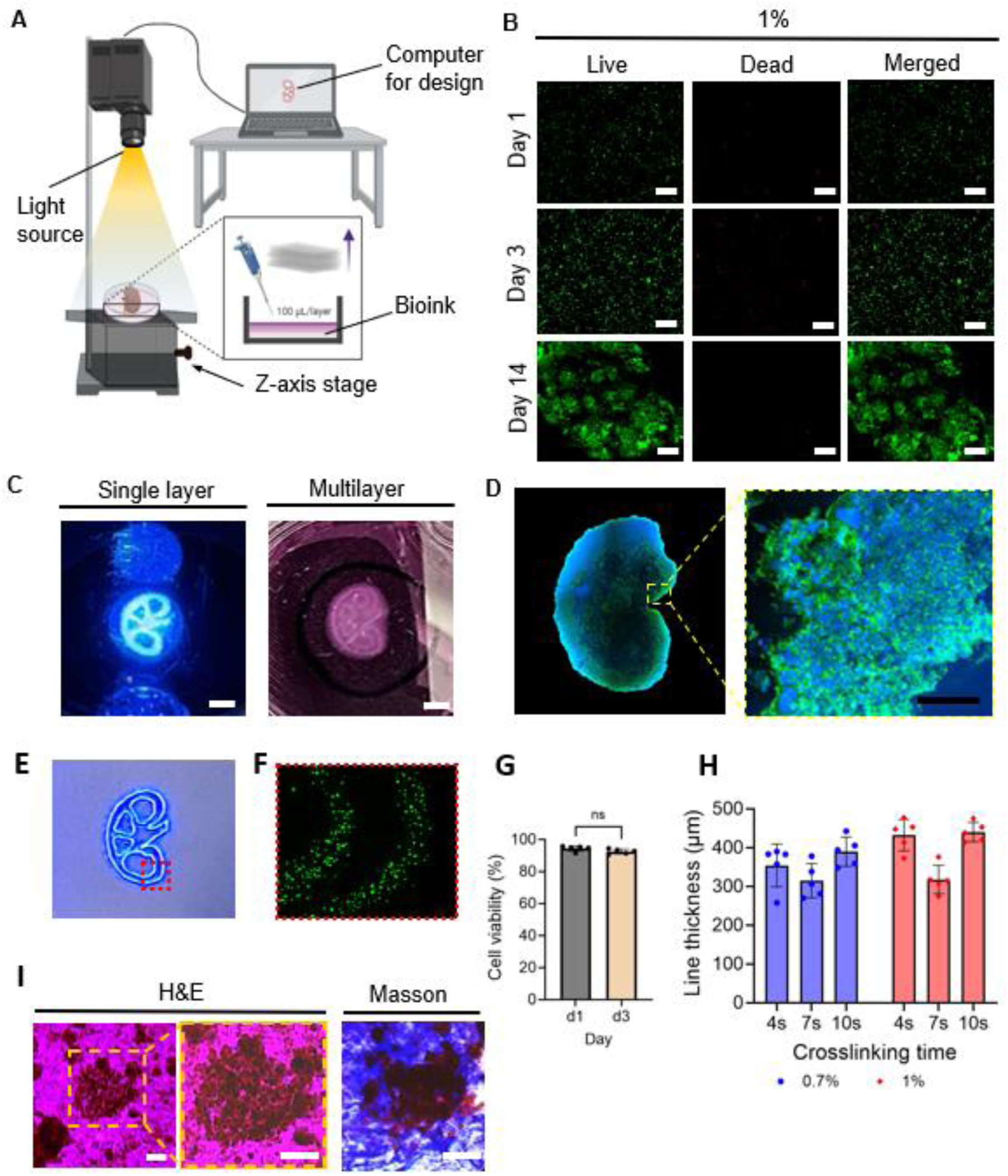
(A) DLP-SLA bioprinting setup equipped with a 405 nm light source and a vertically moving stage, enabling multilayer bioprinting. (B) Representative images of live and dead HEK cells encapsulated in hydrogels over 14 days of culture, Scale bar, 100 μm. (C) The left image shows single-layer printing after 4-second crosslinking with 1% KdMA, while the right image presents an intermediate intersection during the printing process, Scale bar, 5 mm. (D) Confocal imaging of a 3D mini kidney organ with multiple layers bioprinted using 1% KdMA. Scale bar, 200 μm. (E) Intersection design in kidney 3D scaffold printing. (F) Clarity image showing cell encapsulation in a line within the intersection design. (G) Cell viability of the 3D bioprinted scaffold (*n*=5). (H) Printed line thickness based on crosslinking time (*n*=5). (I) Histological evaluation of bioprinted structures. H&E and Masson trichrome staining of a 0.7% KdMA sample after 14 days of HEK cell culture. Scale bars, 50 μm

Structural integrity and multilayer assembly of the bioprinted scaffolds were further analyzed in **Figure 4C**, where the left panel illustrates a single-layer construct, while the right panel presents the multilayer bioprinting process captured at an intermediate stage. As the layers were stacked, the intersections in the scaffold design became less distinguishable due to increased optical density and structural complexity. To better visualize the scaffold architecture, the image was captured when approximately half of the total layers had been printed. To evaluate long-term cellular organization, the bioprinted scaffolds were cultured for two weeks, followed by confocal microscopy imaging. Phalloidin and DAPI staining were performed to visualize cell proliferation and structural organization, confirming that the bioprinted constructs facilitated the formation of a 3D mini kidney-like structure, demonstrating the potential of the developed bioink for kidney tissue engineering applications (**Figure 4D**).

#### 3.5.2. Evaluation of the printability of the deveopled bioink in extrusion-based bioprinting

To further evaluate the bioprinting capability of the developed bioink, an in-house modified extrusion-based bioprinter was utilized (**Figure 5A**). A customized piston-driven extrusion system was implemented, with a stepper motor directly installed, and key components of the bioprinter were 3D-printed to enable precise extrusion control. To facilitate high-cell-concentration bioprinting, a 500 µL syringe was mounted, allowing for the bioprinting of cellular droplets and a 2×2 mesh pattern, while simultaneously crosslinking the printed structures using 405 nm light (**Figures 5B**). As shown in **Figure 5C** (top left), cells within the bioprinted droplets were observed immediately after printing, and fluorescence microscopy imaging confirmed that the cells exhibited elongation after one week of culture (**Figure 5C**, top right), indicating active cell spreading and adaptation within the printed environment. Additionally, the mesh-patterned bioprinted scaffold supported cell proliferation and growth, further demonstrating the versatility and biocompatibility of the developed bioink (**Figure 5C**, bottom).

**Figure 5.**
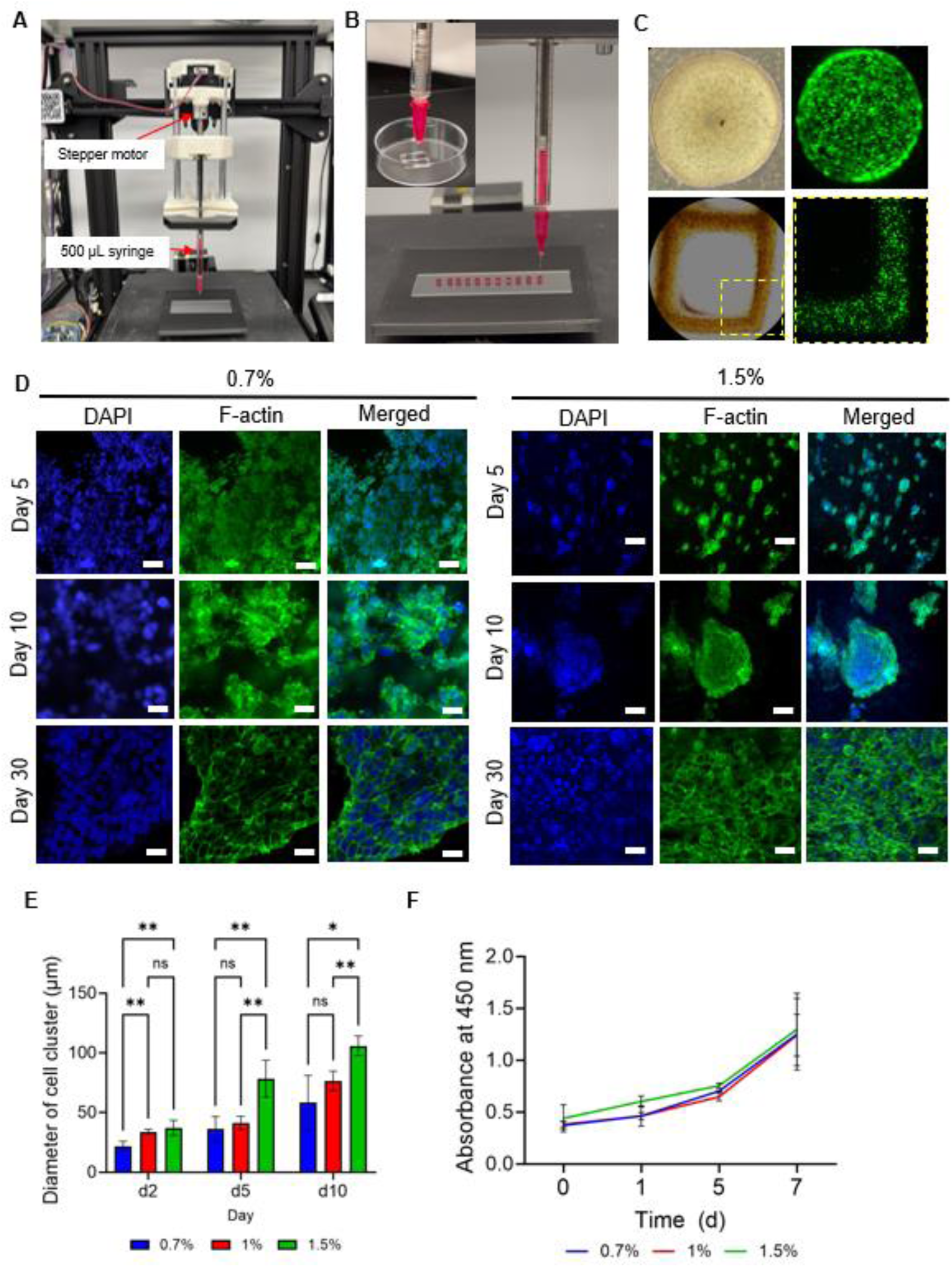
(A) Customized piston-driven extrusion bioprinter setup. (B) Bioprinting of lattice patterns and cellular droplets. (C) Crosslinked cellular scaffolds after bioprinting. Bright-field imaging (top left) shows cell distribution immediately after printing, while flurescence imaging (top right) after one week of culture shows homogeneous cell elongation within the droplets; a magnified view of the lower square region of the lattice pattern (bottom left) and cell viability assessment after printing (bottom right) are also presented. (D) Evaluation of cell growth, proliferation, and morphology over 30 days of culture. Fluorescence images showing HEK cells morphology during the culture period, with phalloidin-stained F-actin (green) and DAPI-stained nuclei (blue), and merged images. Scale bars, 100 μm. (E) HEK cells cluster size distribution at different time points during culture (*n*=5). (F) XTT assay showing HEK cell metabolism and proliferation over 7 days of culture within a 3D hydrogel structure (*n*=5).

To investigate cellular morphology and organizational patterns within the bioprinted hydrogel scaffold, cell cytoskeleton and nuclei staining were performed. The cultures were incubated for one month, and the study demonstrated the progressive development of HEK cells, transitioning from individual cells to cohesive clusters, which ultimately formed multicellular spheroids, similar to previous studies [30]. These spheroids were viable and stable throughout the study period, maintaining a uniform distribution across the entire 3D scaffold up to day 10. In samples with a higher concentration of 1.5% KdMA, cluster size was significantly larger on days 5 and 10 compared to the 0.7% KdMA samples, indicating enhanced cell proliferation (**Figure 5D**). These clusters were characterized by well-defined cytoskeletons and exhibited growth into larger multicellular aggregates over the culture period. Interestingly, by day 30, all tested KdMA concentration samples exhibited multiple proliferating and merging spheroids, demonstrating that the ECM proteins and growth factors inherent in the dECM components within the scaffold significantly promoted cellular diffusion and proliferation.

During cell growth within the bioprinted scaffold, the diameter of cell clusters was continuously tracked. In all bioink formulations, HEK cell cluster size increased with longer culture duration (**Figure 5E**). Notably, the highest bioink concentration of 1.5% KdMA resulted in the largest cell clusters. To assess cell metabolic activity, an XTT assay was performed, demonstrating that the bioink developed in this study supported the HEK cell proliferation (**Figure 5F**). Among the tested conditions, the 1.5% KdMA formulation exhibited the steepest growth rate within the first five days of culture.

### 3.6. Histology assessment

This characterization was conducted to assess the histological differences observed in HEK cells cultured on KdMA scaffolds (**Figure 5G**). To highlight cellular morphology and interactions with the substrate, a standardized H&E staining protocol was applied to 0.7% KdMA scaffolds cultured for 14 days. In H&E-stained samples, HEK cells exhibited characteristic cytoplasmic staining, with nuclei clearly delineated in vivid blue due to hematoxylin staining, while the cytoplasm appeared bright pink from eosin staining. This interaction suggests a specific affinity between eosin—an acidic dye that preferentially binds to cytoplasmic protein components—and the basic cellular structures. The ECM is known to play a crucial role in cellular function and gene expression regulation, influencing intracellular protein synthesis [31]. As a result, the intense cytoplasmic staining observed in HEK cells on KdMA scaffolds suggests robust cytoplasmic protein synthesis, potentially indicating enhanced cellular maturation. Additionally, Masson’s trichrome staining was used to confirm the presence of collagen fibers within the scaffolds supporting the HEK cell culture. The 0.7% KdMA samples demonstrated that the intrinsic collagen fibers in KdMA stabilized through covalent crosslinking, and remained structurally intact and resistant to degradation even after two weeks of culture. This finding highlights the long-term stability of the scaffold, reinforcing the advantage of KdMA bioink for extended kidney cell culture.

## 4. Conclusion

In conclusion, this study successfully demonstrates the development and application of a stiffness-tunable, photocrosslinkable, kidney-specific ECM-based bioink for kidney tissue and organoid bioprinting. The decellularization process effectively removed all cellular components while preserving key structural and biochemical features of the kidney extracellular matrix, ensuring an organ-specific microenvironment that supports cellular growth and maturation. The resulting dECM-based bioink exhibited favorable rheological properties, making it well-suited for both DLP-SLA bioprinting and piston-driven extrusion bioprinting. Furthermore, HEK cells encapsulated within this bioink maintained high viability and proliferative capacity, demonstrating the potential for tissue-specific maturation over time. Notably, this bioink created a kidney-specific microenvironment that actively supported HEK cell maturation and tissue formation, highlighting its potential to enhance organoid maturation. The bioprinted kidney tissue constructs exhibited high cell viability and proliferation, further demonstrating the bioink’s ability to support functional tissue development.

Moving forward, 3D bioprinting strategies that utilize kidney dECM-based bioinks hold significant promise for advancing bioengineered kidney tissue constructs, with implications for regenerative medicine and precision medicine. By enabling the fabrication of engineered kidney tissues, this approach could facilitate therapeutic applications such as drug testing, and disease modeling, and potentially offer alternative solutions for kidney transplantation. Ongoing research is currently investigating how integrating bioinks with iPSC-derived organoid bioprinting affects their interactions and optimizes their potential for clinical translation.

## Acknowledgment

This work was supported by a Natural Sciences and Engineering Research Council of Canada (NSERC) Discovery Grant (RGPIN-2020-04559) and the Canada Foundation for Innovation John R. Evans Leaders Opportunity Fund. In addition, J.S. was supported by the NSERC Vanier Canada Graduate Scholarship and Alberta Innovate Graduate Student Scholarship.

